# Site-specific Crosslinking Coupled with Mass Spectrometry as a Structural Tool in Studies of the Human α_1_ Glycine Receptor

**DOI:** 10.1101/2020.06.29.178681

**Authors:** Rathna J. Veeramachaneni, Chelsee A. Donelan, Kayce A. Tomcho, Shaili Aggarwal, David J. Lapinsky, Michael Cascio

**Author notes:** **Corresponding Author**, • Michael Cascio.

## Abstract

Recent advances in mass spectrometry coupled with chemical crosslinking (CX-MS) can be applied for the structural interrogation of macromolecular complexes to identify statedependent distance constraints and provides a powerful complementary technique to other structural methods. In this study, we develop a CX-MS approach to identify the sites of crosslinking from a single targeted location within the human glycine receptor (GlyR) in a single apo/resting state. The GlyR belongs to the family of pentameric ligand-gated ion channel receptors that function in fast neuronal transmission. A single cysteine residue was re-introduced into *Cys null* GlyR construct at position 41 within the extracellular domain of an overexpressed human a1 homomeric GlyR. After purification and reconstitution into vesicles, a methanethiosulfonate benzophenone heterobifunctional crosslinker was attached via formation of a disulfide bond, and the resting receptor is subsequently photocrosslinked. Monomeric and oligomeric bands from SDS-PAGE gels were then trypsinized and analyzed by tandem MS in bottom-up studies. Dozens of intra- and inter-subunit sites of crosslinking were differentiated and identified from single gel bands (pmols of purified GlyR), showing the utility of this approach to identify a diverse array of distance constraints of GlyR in its resting state. These studies highlight the potential of CX-MS as an experimental approach to identify state-dependent crosslinks of full length integral membrane protein assemblies in a native-like lipid environment.

## Introduction

Despite advances in high resolution structural methods such as X-ray crystallography, NMR, and cryo-EM, membrane proteins remain relatively underrepresented in the PDB database. These methods provide invaluable information on the structure of membrane proteins, but they are not without limitations or difficulties. For example, the presence of the bilayer provides a barrier to both crystallographic and NMR studies. Additionally, X-ray crystallographic and cryo-EM studies provide information on static images of protein that is often stabilized in a single low-energy state. Thus, determination of high-resolution structures is often accomplished by truncating the more flexible regions of the protein, as well as by mutating some residues for stability/homogeneity and using other proteins, such as antibodies and/or ligands, to non-covalently bind and stabilize the protein. Although NMR can be used to study the flexible regions of proteins, this technique is efficient for small-to medium-sized proteins. In large proteins significant spectral overlap occurs, requiring higher field strengths and/or isotopic labeling to allow formal assignment of the spectra. Also, the presence of a lipid bilayer or large micelles also introduces problems as these complexes move very slowly in solution, unlike smaller soluble proteins with higher tumbling rates that are essential for an effective relaxation of the molecule^1^.

For pentameric ligand-gated ion channels (pLGICs), crystallography has provided snapshots of many of these ion channels in ligand-bound forms^2–10^, though the large intracellular loops connecting the transmembrane helices remain poorly resolved. CryoEM is an powerful emerging technique to obtain structural information of membrane proteins in their native state but accurately determining the center and orientation of each image and complications due to sample heterogeneity and the dynamic nature of proteins are potential drawbacks^11^. This methodology provided some of our first images of nicotinic acetylcholine receptors^12,13^ and recent technological advances have provided the field with a wealth of recent high-resolution structures^14–19^, though these often lack information regarding the large intracellular loops and are conducted in a membrane-mimetic environment rather than native-like lipid bilayers. The development of novel experimental approaches that can directly examine mammalian proteins in a native lipid environment and provide inputs to validate and refine computational models can be transformative in allowing validation and refinement of these models.

Given the recent advances in MS technology and its exquisite sensitivity, crosslinking coupled with mass spectrometry (CXMS), has the potential to be a powerful complementary structural tool^20–22^. Importantly, this approach can be used under conditions that challenge or limit other biophysical techniques. CXMS studies do not require large quantities of protein and are not precluded by the presence of a lipid bilayer, so are amenable for studying proteins embedded in lipids. Studies can also be conducted under various physiological conditions (e.g., in presence or absence of agonists, antagonists, or modulatory ligands) to potentially probe state-dependent allosteric changes in large oligomeric complexes. Importantly, the exquisite sensitivity of modern mass spectrometers allows one to examine proteins that can only be isolated at the relatively low concentrations typical of membrane protein preparations.

Chemical crosslinking using bifunctional crosslinking agents can provide distance constraints between the reactive sites within a protein, as defined by the length of the crosslinker. Typical chemistries employed with bifunctional crosslinkers are amine-reactive crosslinkers that make use of abundantly available lysine residues in the protein^23,24^, thiol-reactive crosslinkers that forms a disulfide linkage with reactive cysteines^25,26^, carboxylate-reactive crosslinkers that activates carboxyl groups for spontaneous peptide-bond formation with primary amines^27^, and non-specific crosslinking using photoreactive functional groups to generate highly reactive radicals that interrogate the local chemical environment.

Once crosslinked, mass-shifts due to the introduced moiety can be identified using MS. A bottom-up approach is typically employed wherein proteins of interest are crosslinked and subjected to enzymatic digestion. Additional information may also be obtained by MS/MS analysis of precursor ions to fragment the peptide further to identify/refine the site of covalent crosslinking within a peptide via analyses of the product mass ions after dissociation by collision induced dissociation (CID), electron transfer dissociation (ETD) or electron capture dissociation (ECD)^28–30^. Identifying the sites of chemical crosslinks generated within a protein complex in a given allosteric state yields a network of crosslinks, whichprovide distance constraints that may be used to define the local protein environment in a state-specific manner.

GlyRs are inhibitory ligand-gated receptors that belong to pLGIC superfamily. GlyRs mediate neurotransmission in the central nervous system and are typically activated by glycine. Inhibitory members of the pLGIC receptor family (also referred to as the Cys-loop superfamily) include GlyR, glutamate-gated chloride channel receptors (GluCIRs) and GABA_A_ receptors. These ionotropic receptors show promising silencing activity^31,32^, as these chloride channels typically allow the passive influx of Cl^-^ that further hyperpolarizes the cell under typical conditions. The receptors in this family are pentamers pseudo-symmetrically arranged in a ring around a central ion-conducting pore. Each subunit consists of an extracellular N-terminal ligand-binding domain (ECD), four transmembrane helices (M1-M4), and an intracellular domain. The intracellular domain is primarily comprised of a long loop of variable length between M3 and M4^33^, which currently remains unresolved in all reported structures of pLGICs.

While GlyR can exist in multiple allosteric states as the channel gates and desensitizes, in the absence of ligand (glycine), the receptor is essentially in a resting state. GlyR can be composed of various combinations of α1, α2, α3, α4 and β subunits. Typically, in adult humans it is composed of two α subunits and three β subunits^34^. However, expression of human α1 subunits is sufficient to reconstitute homomeric receptors with native-like function^35^. In this study we develop a novel methodology to conduct protein-protein crosslinking studies of full-length homomeric α1 glycine receptor (GlyR) reconstituted in lipid vesicles in order to identify intra- and intersubunit crosslinks in the resting (apo) state of the receptor. For this work, human α1 GlyR homopentamers, containing a single reactive thiol at Cys41 in each subunit, were overexpressed in insect cells using a baculovirus expression system, then purified and reconstituted using established methods^35,36^. There are 7 endogenous Cys in α1 subunits, four of which form two essential disulfide bonds, leaving three Cys with free reactive nucleophilic thiols. These three cysteine residues have been mutated to provide a *Cys null* background to allow for the systematic introduction of single Cys in crosslinking/MS studies. This engineered single reactive thiol within each subunit was then selectively targeted by MTS-benzophenone (MTS-bzp), a heterobifunctional crosslinker that is initially bound by a disulfide linkage to the mutated site when added to the purified, reconstituted GlyR. The bzp moiety was subsequently photoactivated to interrogate the local protein structure. Intra- and inter-subunit crosslinks were identified by bottom up MS analysis of trypsinized peptide fragments from monomeric and oligomeric bands, respectively, in SDS-PAGE gels. The electrospray ionization-time of flight-mass spectrometry fitted with a chipcube (chip-nESI-TOF-MS) was used to analyze mass-shifted tryptic peptides. Analysis of the pattern of daughter ions, allowed for precise identification of site(s) of crosslinking within the parent peptide. The development of a bottom-up approach using CXMS provides an invaluable experimental approach in providing distance constraints between the residues in the resting receptor and demonstrates the capability of CXMS to sensitively and accurately probe the structure of the resting state of GlyR in an intact lipid bilayer.

## Experimental Section

### GlyR mutagenesis, purification and reconstitution

A *Cys null* construct of the human α1 GlyR (a C41A/C290S/C345A triple mutant in which three reactive Cys present in wild type α1 are mutated to Ala or Ser residues) in pFastBac was used as a template for site directed mutagenesis to re-engineer an endogenous Cys back at position 41, resulting in a mutated form of the receptor that only has a single reactive Cys per subunit. This mutation was introduced using Quick-change site-directed mutagenesis kit (Stratagene) following the manufacturer’s protocols^37^ The forward primer had the sequence CCAGTGAACGTGAGC<u>TGC</u>AACATTTTCATCAAC (the codon for Cys is underlined). PCR conditions used were 16 cycles of denaturation at 95 °C for 30 sec, annealing at 55 °C for 1 min and an elongation at 68 °C for 7 min. Parental plasmid after PCR were cleaved by Dpn digestion and the reaction product were transformed into 50 μL of XL-1 blue supercompetent cells by heat shock treatment at 42°C for 45 sec. Cells were allowed to recover in NZY^+^ broth while shaking for an hour at 37°C and then plated on LB (Luria Bertani) agar plates containing 50μg/mL ampicillin. After overnight incubation at 37°C, single colonies were isolated. Plasmids were extracted and purified using a Bio-Rad mini prep kit, and the correct mutation was confirmed by sequencing (GENEWIZ).

Production of recombinant baculovirus followed the manufacturer’s protocol (Invitrogen). Briefly, the purified mutated pFastbac construct was transformed into MAX Efficiency DH10Bac™ competent cells by heat shock treatment for 105 sec at 42°C, and then grown in Super Optimal broth with Catabolite repression (S.O.C) media by incubating it at 37°C for 4 hours. 100 μl of serial dilutions of 10^-1^, 10^-2^, 10^-3^ samples were plated on Luria agar plates containing kanamycin (50 μg/mL), tetracycline (10 μg/mL), and gentamycin (7 μg/mL), Bluo-gal (300 μg/mL) and IPTG (40 μg/mL) and incubated for 24 to 48 hours at 37°C. The presence of white colonies amidst the blue colonies indicates the presence of recombinant bacmid DNA. White colonies were picked and grown in LB media containing antibiotics at 37°C for 24 hours and the recombinant bacmid DNA isolated. Recombinant DNA (1-2 μg) was transfected into *Sf-9* (*Spodoptera frugiperda*) insect cells (cell density at 5 x 10^5^ cells/mL) using 6 μL of CellFectin reagent and 100μL of SFM II media with antibiotics (penstrep, 0.5X) (Gibco™) and allowed to grow for 1 hour at 28°C. The supernatant was then replaced with SFM II media without antibiotics and grown for 72 hours at 28°C. After viral amplification of the harvested supernatant, the plaque forming units (pfu)/mL was determined by viral titer. Briefly, 25 uL of serially diluted (10^-2^ – 10^-6^) virus were added in duplicates to insect cells seeded wells containing 3-5 x 10^5^ cells/mL and allowed to bind for 1 hour. The virus was aspirated, and cells washed with PBS buffer, pH 7. Cells were fixed using 10% gelatin for 10 minutes. A primary antibody mouse anti-gp64 (a glycoprotein expressed on the surface of infected cells) (Abcam)^38^ was added to the plate (1:200), incubated for 1 hour at 37°C, washed twice with PBS buffer (pH7.2) and developed by incubation with goat anti-mouse Ab-HRP conjugate (Abcam) (1:1000) and incubated for 1 hour at 37°C. After washing twice, TMB substrate (ThermoFisher Scientific) was added and the plate was incubated at room temperature for 30 minutes. The number of blue stained cells per well were counted and the number of cells and the dilution factor was taken into consideration to calculate the pfu/mL. The viral particles were further used to infect cells in higher quantities for overexpression of the mutated protein^39^. The tittered virus is then added to a spinner flask (volume determined based on using a multiplicity of infection of ~ 5) containing about 600 mL of cells at a concentration of ~1×10^6^ cells/mL and >97% viability. The cells were harvested 3 days post-infection^36^.

### GlyR purification and reconstitution

GlyR purification and reconstitution were performed as previously described^36^. Briefly, the cell pellet was re-suspended and washed three times in an isotonic phosphate buffered saline solution. The cells were suspended in a hypotonic solution containing 5 mM Tris (pH 8.0), 5 mM EDTA, 5 mM EGTA, and 10 mM dithiothreitol for 1 hour at 4 °C to swell the cells prior to lysing. An anti-proteolytic cocktail containing 16 milliunits/mL aprotinin, l μM benzamidine, l μM phenylmethylsulfonyl fluoride, and 1 μM benzethonium chloride is added immediately before sonication using a probe tip sonicator with a micro tip attachment (Branson Ultrasonics Sonifier Model 250) using a 50% duty cycles (1/2 sec bursts each second for 15 times) in an ice jacketed container to minimize sample heating. Lysates are centrifuged at 150,000 x g for 30 min. The pellet was re-suspended in a hypertonic saline solution (hypotonic solution + 300mM NaCl) sonicated, and re-pelleted to extract peripheral membrane proteins. The pellet was incubated in a solubilization buffer containing 1% digitonin:0.1% deoxycholate, 1.5 mg/mL mixed lipids (9:1 plant extract (Avanti^®^):egg extract (Avanti^®^)), 25 mM potassium phosphate, pH 7.4, 1 M potassium chloride, 5 mM EDTA, 5 mM EGTA, 10 mM DTT and the anti-proteolytic cocktail by sonication and rocking for 48 hours. Samples were centrifuged as before, and the supernatant was added to 2-aminostrychnine agarose resin and, incubated for 24 hours at 4°C with nutation. After extensive washing with PBS (pH 7.4), bound lipid/protein/detergent micelles were eluted by 2-day equilibration in solubilization buffer with added 200mM glycine. Purified protein-lipid vesicles were reconstituted by dialysis (3500 Da cutoff Slide-A-Lyzer dialysis cassette, ThermoFisher Scientific) against excess 25mM phosphate buffer (pH 7.4) over 5 days. Post-dialysis, the reconstituted GlyR is sonicated as before and pelleted at 150,000 x g for 30 min to concentrate the vesicles that are then re-suspended in 1 mL of 25mM potassium phosphate.

A modified Lowry assay^40^ was used to determine the concentration of GlyR. For quality control, aliquots from various steps of the purification were visualized by SDS-PAGE and Western blotting. Primary antibody (rabbit anti-GlyR, Abcam) at 25,000 dilution and IR-labeled secondary antibody (goat anti-rabbit, LI-COR) at 10,000x dilution were used. Membranes were visualized using an ODYSSEY CLx infrared imager (Li-Cor).

### Chemical synthesis of S-(2-(4-(4-(prop-2-yn-1-yloxy)benzoyl)phenoxy)ethyl) methanesulfonothioate (MTS-bzp)

Sodium methanethiosulfonate (75 mg, 0.56 mmol, 1.4 eq.) was added to a solution of (4-(2-bromoethoxy)phenyl)(4-(prop-2-yn-1-yloxy)phenyl)methanone^41^ (140 mg, 0.39 mmol, 1.0 eq.) in DMF (9.3 mL) at room temperature. The resulting mixture was then heated at 50°C for 20 hours under argon, wherein TLC analysis in 3:7 EtOAc:hexanes indicated a complete reaction. The reaction mixture was then cooled to room temperature, diluted with H_2_O, and extracted with three times with EtOAc. The combined EtOAc extracts were then dried (MgSO_4_), filtered, and concentrated in vacuo to give 150 mg (99%) of the target MTS-benzophenone-alkyne as a sticky yellow solid requiring no further purification. Rf = 0.61 (95:5 CHCl_3_:MeOH). ^1^H NMR (400 MHz, Chloroform-*d*) δ 7.78 (d, *J* = 8.8 Hz, 4H), 7.04 (d, *J* = 8.8 Hz, 2H), 6.96 (d, *J* = 8.8 Hz, 2H), 4.77 (d, *J* = 2.5 Hz, 2H), 4.37 (t, *J* = 6.0 Hz, 2H), 3.58 (t, *J* = 5.9 Hz, 2H), 3.42 (s, 3H), 2.58 (t, *J* = 2.4 Hz, 1H). ^13^C NMR (101 MHz, Chloroform-*d*) δ 194.1, 161.1, 160.6, 132.2, 132.0, 131.3, 131.2, 114.3, 113.9, 77.8, 76.1, 66.7, 55.8, 50.7, 35.4. HRMS calculated for (C_19_H_18_O_5_S_2_)Na^+^ 413.0488, found 413.0493.

### Crosslinking

250 μL aliquots of GlyR vesicles in 25 mM phosphate buffer (pH 7.5) and MTS-bzp (1.9 mg/mL in DMSO) were combined to give a 1:5 GlyR:MTS-bzp mole ratio (one crosslinker per subunit of pentameric GlyR) and allowed to react overnight in the dark while nutating. The sample was transferred to a quartz cuvette and exposed to a 420 W Hg Arc lamp (Newport, Model 97435-1000-1, 260-320 nm) for 4 sessions of 5 minutes on ice, with 5 minute periods of no exposure in between each UV exposure session to prevent sample warming. 20 μL of crosslinked samples were loaded onto a 15% SDS-PAGE as described previously except under non-reducing conditions (no dithiothreitol) in order to be able to separate α1 subunits containing potential intersubunit crosslinks (oligomeric bands) from those containing solely intrasubunit crosslinks (monomeric bands). Given the low concentrations of GlyR used in each trial, marker lanes containing bovine serum albumin (MW 66.5 kDa) (Sigma-Aldrich) were used to direct gel excision. Gel plugs corresponding to low MW monomeric (intrasubunit crosslinks) and high MW oligomeric (inter- and intrasubunit crosslinks) were excised.

### Trypsinolysis and MS sample preparation

Gel plugs were washed with 50:50 methanol: 50mM ammonium bicarbonate solution and then dried with acetonitrile. The protein in the gel plugs was reduced using 10 mM DTT and alkylated with 50 mM iodoacetamide. Gel plugs were soaked in 50 mM ammonium bicarbonate (pH 7.8) containing 20 μg/mL of trypsin gold (Promega) for 24 hours at 37°C. Digested peptides in the supernatant from the gel plug were transferred into a fresh tube. 0.1% formic acid (FA) in 50:50 acetonitrile (ACN): H_2_O is added to gel plugs incubated for 30 minutes and the supernatant is transferred into the initial elution to deactivate the trypsin enzyme. The extracted solution is evaporated and is re-constituted in 20 μL of 0.1% FA in Milli-Q^®^ water. While SDS-PAGE gel denatures the globular structure of GlyR, trypsin digests the GlyR subunits into peptides for MS to detect. Peptide samples are further cleaned up and concentrated using Zip Tips (EMD Millipore). Zip Tips are 10 μL pipette tips with a 0.6 μL bed embedded with a very small, reverse-phase C18 resin. These Zip Tips are conditioned using 50% ACN three times and equilibrated with 0.1% FA three times. The peptide sample is then pipetted through the column 10-20 times to allow the peptides in the sample to bind to the column. The column is then washed with 0.1% FA to remove any impurities and the peptides bound to the column are eluted using 10 μL of 0.1% FA in 50% methanol.

### Mass spectrometry

Samples were run on an Agilent accurate Mass 6530 nano-electrospray ionization time-of-flight mass spectrometer (nESI-QTOF-MS) fitted with a C18 reverse phase (75 μm X 150mm separation column and a 40 nL enrichment column) chip cube. The mass spectrometer was run in positive ion mode using internal standards (1221.9906 and 299.2944) for calibration, supplied by Agilent. Samples were re-suspended in 40 μL of 0.1% formic acid in water. Each run was conducted using a 2μL injection volume and a linear gradient using solvents A (95% H_2_O; 5% ACN; 0.1% formic acid) and B (95% ACN; 5% H_2_O; 0.1% formic acid) run over 11 minutes with the profile: 0 min 95% A; 7 min 40% A; 9 min 10% A and 9.10 min 95% A. M/z of 200-1700 and 175 V fragmentor voltage. Qualitative analysis software (Agilent) identified mass-shifted crosslinked precursor mass ions using a 10 ppm cutoff. The crosslinker mass shift of MTS-bzp (311.3838 Da) along with other possible modifications (i.e., sodiation, postassiation, alkylation, oxidation and acrylamidation) were used in identifying the mass shifted precursor ions. For targeted MS/MS by CID, a linear increase in collision energy by m/z was calculated using the equation y= 3.7x + 2.5 (y = m/z, x = collision energy). Data analysis was performed using a 0.1 Da error in product ions using Masshunter (Agilent), Prospector (MS-product UCSF), SkyLine (MACCOSS LABS), and Spectrum Mill (Agilent).

## Results and Discussion

MTS-bzp containing a clickable alkyne tag (**Fig. 1**) was used in crosslinking studies followed by bottom-up LC-MS/MS analysis to identify sites of inter- and intrasubunit covalent attachment. A single Cys was re-introduced into a human a1 *Cys null* GlyR background via mutation of Ala at position 41 (A41C). The mutation at position 41 was selected for these proof-of-principle CX-MS studies as it is an endogenous Cys in the wild type receptor (it was mutated to create the *Cys* null construct) and is known to be accessible from previous studies in our lab. Homopentameric a1 GlyR was overexpressed in *Sf*9 insect cells using a baculovirus expression system, and whole cell patch clamping studies of infected insect cells showed the receptor retained native like activity (data not shown). GlyR was purified and reconstituted into vesicles containing cholesterol and mixed lipids from plant and chicken eggs using established methodologies that reconstitute GlyR in a fully functional form^42^.

**Fig. 1.**
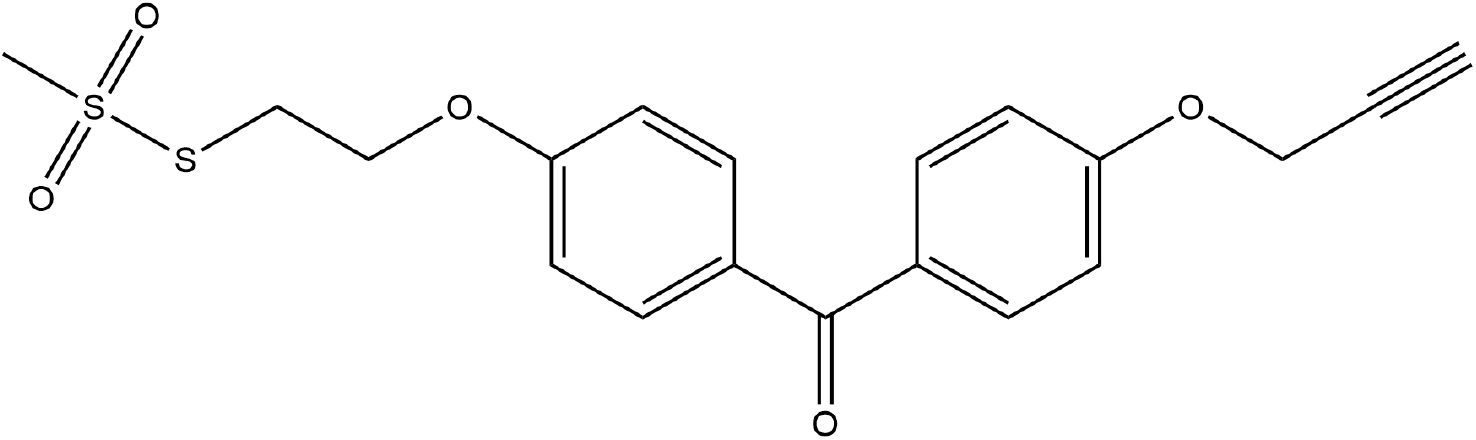
MTS-BzP crosslinker. Thiol reactive MTS group is shown on left, photoreactive benzophenone is in center and clickable alkyne is on right.

The heterobifunctional crosslinker MTS-bzp (**Fig. 1**) contains an MTS moiety that covalently binds to the sulfhydryl group of single engineered Cys in each subunit of the human homopentameric a1 GlyR, providing an initial site of attachment that is sitespecific. The crosslinking agent contains a photoactivatable benzophenone (bzp) moiety that preferentially targets local C_α_ backbone atoms upon photoactivation^43^ and a terminal alkyne tag that can be further derivatized using click chemistry^44^. Upon photoactivation bzp has been reported to undergo either a H-abstraction or a H-abstraction followed by a recombination mechanism^43^ (**Fig. 2)**. These mechanisms can be differentiated by MS, as the resultant masses will differ by the mass of a H atom and may be distinguished given the sensitivity of MS platforms (we utilize a 10 ppm error window in m/z assignments). The observed crosslinking products are consistent with the latter H-abstraction and recombination mechanism (**Fig. 2**, bottom right).

**Fig. 2.**
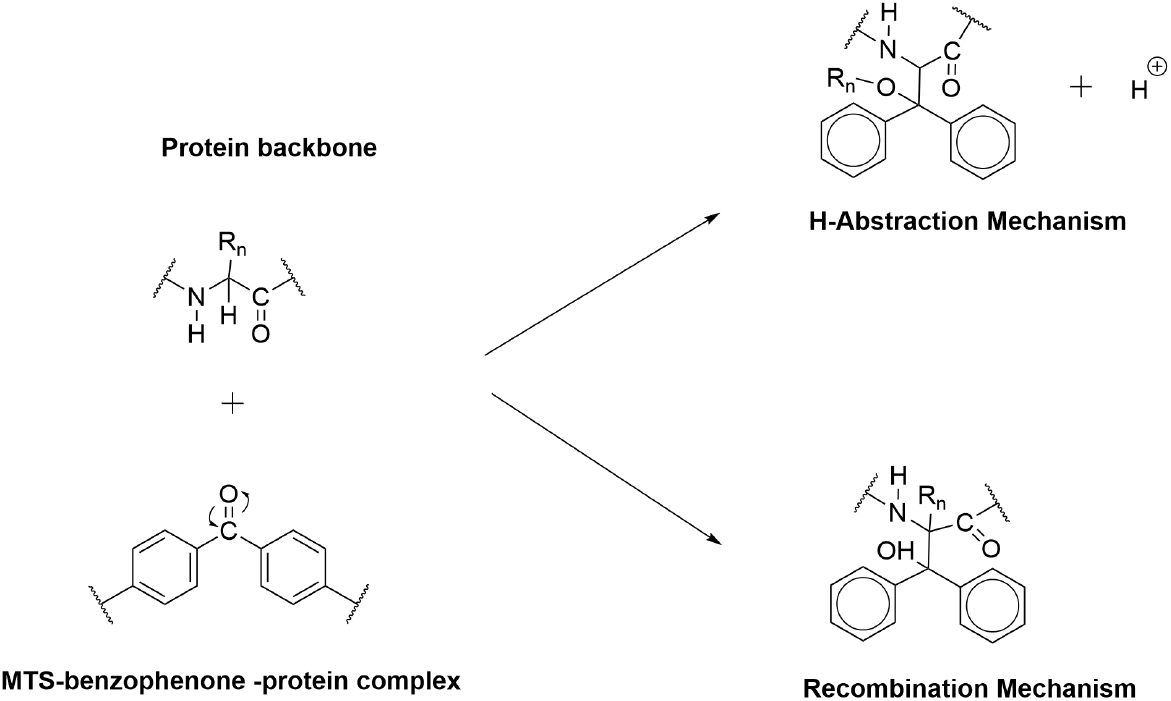
Proposed benzophenone photocrosslinking mechanisms.

SDS-PAGE separation in the absence of reductant was used to separate oligomers from monomers. Similar to other membrane proteins, GlyR runs as both monomers and oligomers on SDS-PAGE even in the absence of any crosslinking. Given this, and the presence of five potential crosslinks per pentamer, the mass-shifted crosslinks identified in oligomeric bands may contain containing either intra- or inter-subunit crosslinks. However, any mass-shifted peptides identified in MS studies of the monomeric band must be intrasubunit crosslinks. The respective excised gel plugs were trypsinized and extracted and then the peptides were reduced and alkylated (separating the site-specific Cys-linked source peptide from photocrosslinked target peptides). Unique mass-shifted peptides observed only in tryptic extracts from the oligomeric band and absent in the extracts from the monomeric band are hypothesized to occur between neighboring subunits. A similar strategy was validated in Lys-Lys crosslinking studies conducted on the extracellular domain of GlyR^45,46^.

Three independent GlyR preparations were separately purified, reconstituted and used in CX-MS studies. Mass-shifted crosslinker-containing peptides with m/z in the 300 – 1200 Da mass range with a mass error of ≤10 ppm were identified. A single missed trypsin cleavage was allowed as were carboxymethylation, Met oxidation, sodiation, potassiation or acrylamide modifications. Identified precursor mass-shifted ions were targeted for MS/MS via fragmentation using collision induced dissociation (CID) using N_2_ gas to confirm their original assignment and to identify the site(s) of crosslinking within the peptide by analyses of the MS/MS spectra.

CID fragmentation typically produces *b* and *y* product ions of the precursor mass ion. Analyses of product ions containing mass shifts due to the attached crosslinker allow refinement of the covalent attachment of the crosslinker, sometimes to a single amino acid site. A representative MS and MS/MS spectra of A41C showing the identified site of crosslinking is shown in **Fig. 3**. Given that tryptic peptides could be modified at more than a single site in a given experiment, MS/MS spectra are strictly matched to their precursor ion by retention time to differentiate isobaric species differing only by the site of crosslinker attachment. For example, multiple amino acids on a given tryptic peptide may be within the non-specific crosslinking radius from Cys41, generating multiple singly-crosslinked species. Each isobaric precursor ion would yield unique patterns of product ions after CID. Given that in-line liquid chromatography can potentially separate isobaric peptides via differential chemical properties, each correlated (with respect to retention time) precursor/product pair can yield a refined crosslinking site(s) even when present at multiple sites. By individually analyzing each product ion fragmentation spectrum at a given retention time, isobars may be unequivocally resolved in MS/MS studies. Only precursor ions assigned to crosslinker-containing peptides (≥ tetrapeptides) identified having overlapping sequence coverage in ≥ 2 of 3 trials are reported in **Table 1**. CID fragmentation within the retention time window (+/- 0.2 minutes) produces product ion fragmentation spectra (2-12 spectra) for each crosslinked peptide. Product ion scans for mass-shifted precursor ions were individually analyzed to identify the site of covalent attachment and the sites of crosslinking within isobaric tryptic fragments are shown in bold in **Table 1**.

**Fig. 3.**
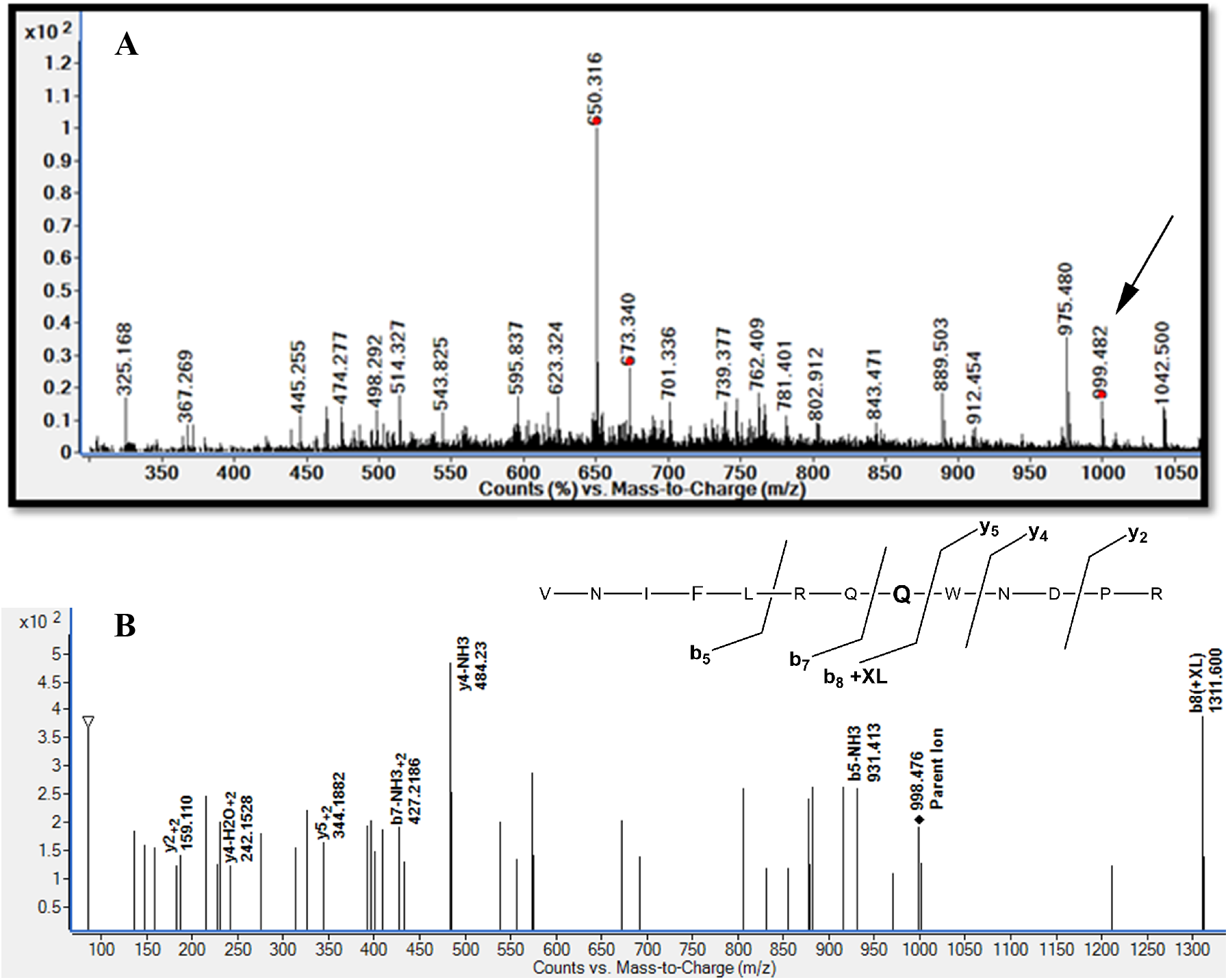
Representative MS-MS analysis. **A.** Representative MS scan identifying crosslinked precursor ion (arrow). **B.** CID-induced fragmentation of crosslinked precursor ion. The identified *b* and *y* product ion fragments, including those mass shifted (+ XL) are identified and refine the site of location within the tryptic peptide (**bold**).

**Table 1.**
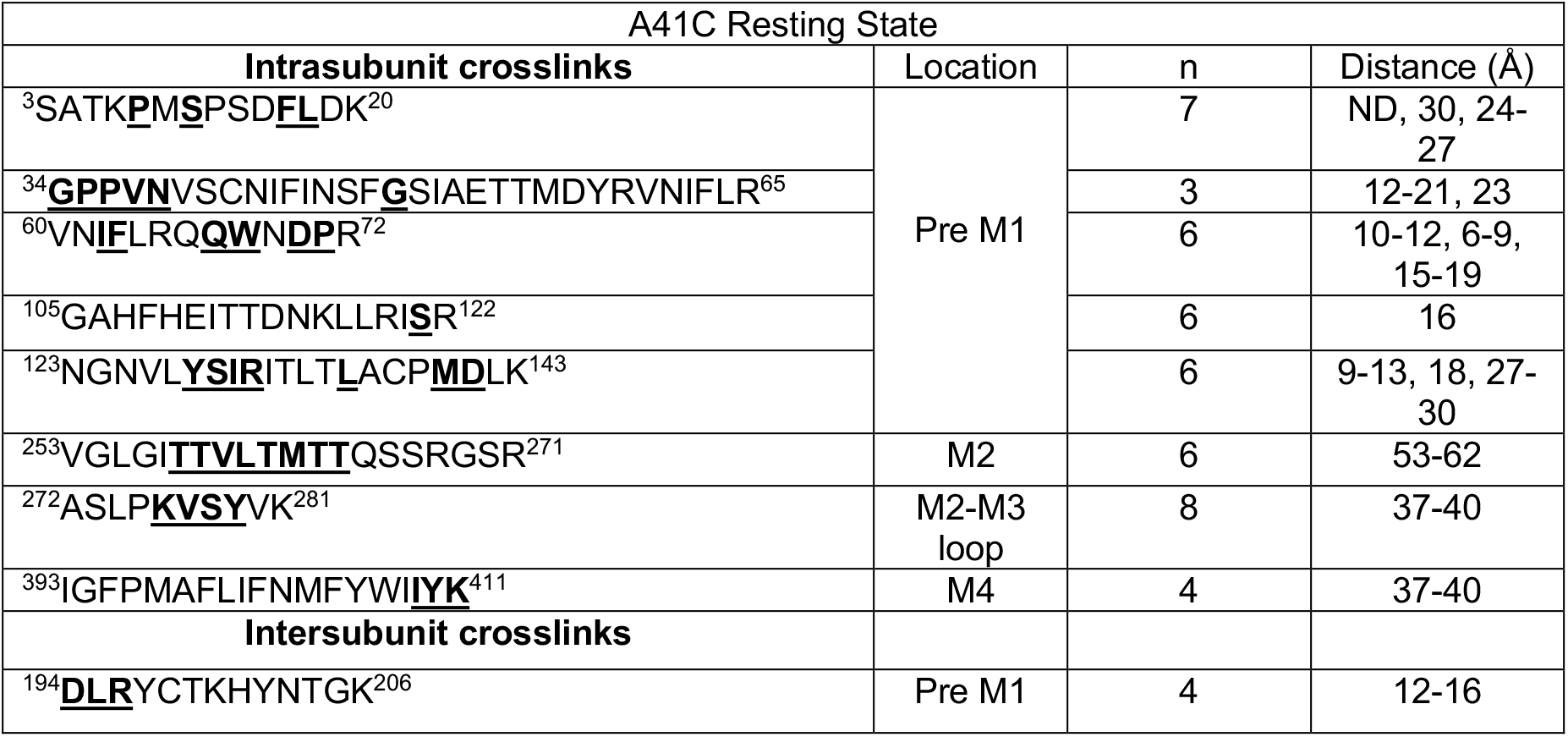
The identified mass shifted tryptic peptides extracted from monomeric and oligomeric bands in SDS-PAGE, with n being the number of trials the peptide was observed within 10 ppm mass difference. The identified crosslinked peptides are further fragmented using CID and unique residue/short peptides identified within 0.1 Da, and the refined site of covalent attachment are shown in **<u>bold and underlined</u>**. The reported distances listed are Cα-Cα distances from C41 to site(s) of identified covalent modification, respectively, in the structure of zebrafish α1 GlyR complexed with strychnine (PDB: 3JAD). Ranges are provided where the site of crosslinking could not be refined to a single amino acid. ND = not determined in crystal structure

All MS runs were carried out in triplicate and the identified crosslinked sites listed were observed in more than one ion species (i.e., protonated, sodiated, potassiated, alkylated and oxidated species). The multiplicity of crosslinking events observed from a single site of attachment, as more than 15 sites of crosslinking each single trial from a single excised SDS-PAGE gel band, shows the utility of using a non-specific photoactivatable crosslinker to interrogate the local protein topography. The promiscuity of binding events provides a large network of distance constraints that can then be used to validate and refine structural models. The introduction of the single site mutation presented here, while consistent with current models of GlyR and pLGICs, is not sufficient to rigorously validate the model. However, we propose that CXMS studies conducted on systematically introduced single Cys mutants will provide a large network of constraints capable of refining the resting state of GlyR, including regions of the receptor that are currently unresolved (e.g., the large intracellular loop connecting the M3 and M4 transmembrane helices. These more comprehensive crosslinking studies will follow this initial proof-of-principle study.

As expected, peptide extracts from higher MW oligomeric bands contained many redundant mass-shifted peptides since oligomers may contain both intra- and intersubunit crosslinks (**Table 1** has been simplified to reduce redundancies in assignments, but a complete list of all assigned crosslinks from each respective gel slice and the respective modifications observed is provided in the Supplementary Information). This redundancy provides further confidence the assignment, and product ion scans from extracts from oligomeric bands provided additional information allowing better refinement of the sites of covalent attachment within the mass-shifted peptide. There was only a single mass-shifted peptide that was reproducibly observed solely in peptide extracts from oligomeric bands. Given this restriction, it is proposed that the bzp moiety attached at C42 forms intersubunit crosslinks to a residue within the range of 194-196 in the pre-M1 region of its neighboring subunit. The intra- and intersubunit residues that the crosslinker bound to are identified for the resting state the single Cys mutants A41C GlyR are shown in **Table 1**. Sites of crosslinking were visualized by mapping the corresponding sites of attachment on a model of the zebrafish α1 GlyR structure (PDB: 3JAD)^19^ (**Fig. 4).**

**Fig. 4.**
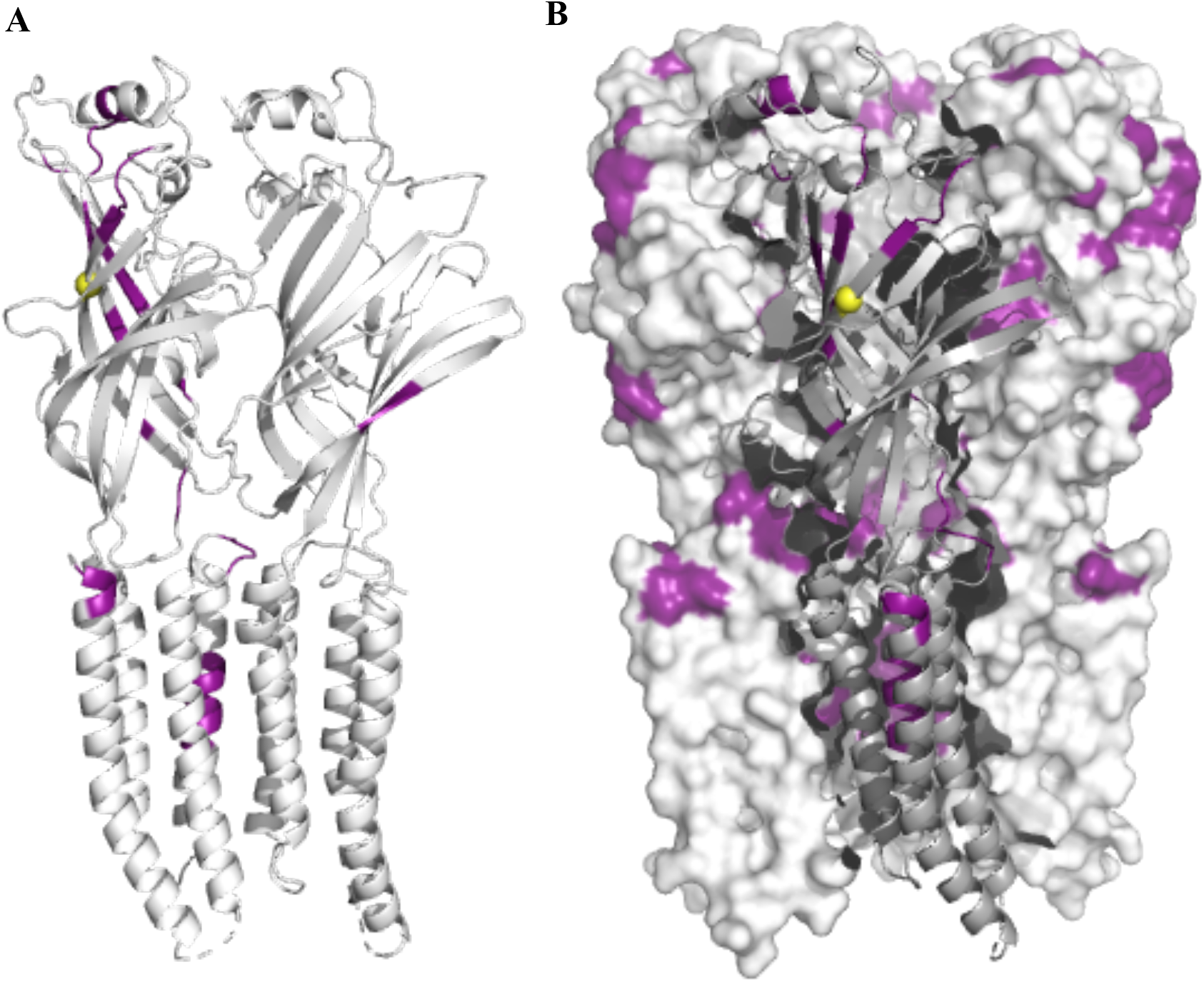
Sites of crosslinking from Cys^41^ mapped on to model of zebrafish GlyR (PDB #3JAD). C_α_ of C41 is shown as yellow ball and crosslinked residues (bold and underlined residues in **Table 1**) are mapped in purple. **A.** Cartoon representation of two neighboring subunits to differentiate intrasubunit crosslinks within (left) and intersubunit crosslinks (right) between subunits. The ECD of GlyR is at top and transmembrane helices embedded in the lipid bilayer are positioned at bottom. Large extracellular loops linking M3 and M4 helices and N and C termini are not shown as they are currently unresolved. **B.** Space filling model of pentameric GlyR with single subunit shown in cartoon form (as in A). Figure in B is partially rotated from that in A for clarity. Figure was made using PyMOL v1.8.

Significantly, as would be expected, the sites of crosslinking from C41 are reproducibly localized to the extracellular domain (ECD), the M2-M3 linker, or the extracellular ends of the TM helices. No crosslinking events were identified in the more cytoplasmic region of the TM helices or to the intracellular domain (these regions comprise a significant portion of the mass of the protein), providing confidence that the observed crosslinks are not artifactual (i.e., due to misfolded protein). One site of crosslinking identified near the N-terminus tail is not indicated on the model, nor is a distance provided, as this flexible region is not resolved, and thus is missing from the PDB file. Information obtained on the N-terminal region of GlyR whose structure is currently unresolved emphasizes the applicability of CX-MS as an orthogonal biophysical technique to complement current high-resolution methodologies.

The C_α_-C_α_ distance of the MTS-benzophenone crosslinker used in these studies is estimated to be ~25 Å. Assuming an upper bound of ~30Å for crosslinking events, the calculated distances from C41 to crosslinked residues in the extracellular domain are consistent with this limit (**Table 1)**. Distances over 30Å are calculated for crosslinks observed to residues identified within the transmembrane regions. However, it should be noted that other investigators have observed systematic conflicts between structural models and *in situ* distance restraints that are attributed to protein dynamics^22,47^. In addition, all distances reported in **Table 1** are measured using a crystal structure of a strychnine-bound form of human GlyR in detergent micelles, as this is the most relevant model currently available. Currently, there is no structure of the apo form of any eukaryotic pLGIC. In addition, it is known that pLGIC structure and function are affected by lipid composition^48–50^, and no structures in native lipid bilayers are currently available. This latter consideration highlights a particular strength of CXMS studies and their potential to complement other high-resolution biophysical methods examining highly allosteric dynamic complexes in a native lipid membrane.

In these studies, milligram quantities of the GlyR were purified in each preparation. Aliquots of these samples were subjected to crosslinking. Given the loading of a typical gel, low microgram quantities of the crosslinked GlyR sample is then applied to SDS-PAGE and subjected to subsequent in-gel trypsin digestion. The eluted sample from a single excised band after proteolysis containing microgram quantities of protein sample is subjected to in-gel trypsin digestion, elution and clean-up steps and typically 1/10th of the eluted peptide solution is injected into the electrospray ion source for MS analysis. At best, if one assumes that initial Cys labeling is complete, and photoactivation always yields in protein-protein crosslinks, then picomoles of peptides in total would be expected to be crosslinked. We identified dozens of crosslinks in each MS run, if these crosslinks are in equal abundance, and an average single run yields > 20 crosslinking events, at most, picomoles of each peptide can be detected. This estimate is clearly high, as yields are not expected to be 100%, nor are the mass-shifted peptides expected to have the same frequency of occurrence. Yet dozens of crosslinks are detected and resolved in MS/MS studies, showing the sensitivity and capability of high-resolution MS in CXMS.

Given initial concerns that low frequency crosslinking events would be difficult to detect amidst the abundance of non-crosslinked tryptic peptides in our extracts, a terminal alkyne tag was included as part of our crosslinker. In initial studies, click chemistry with biotin azide was used to selectively biotinylate only those peptides containing covalently-bound crosslinkers. After trypsin digestion, soft avidin affinity chromatography was used to isolate only the biotinylated crosslinked peptides. It was anticipated this would allow for facile selective purification and post-column enrichment of only the crosslinked peptides in a small elution volume (providing postcolumn concentration of crosslinked peptides for subsequent MS studies). However, prolonged and extensive studies of the biotinylated peptides yielded only negative results in our hands, as no mass-shifted peptides were observed in MS studies either after or before avidin purification. We hypothesize that the biotinylation resulted in ion suppression and failure to identify crosslinked peptides in MS studies. The salts present in buffers used to isolate biotinylated crosslinked peptides from a streptavidin column via affinity enrichment could also be a potential cause of ion suppression. Surprisingly, only when the click chemistry step was omitted was the presence of mass-shifted peptides was detected in MS studies. Developing a clean-up protocol using a solid phase extraction to remove the unwanted salts might be a solution to this problem. However, given our success in identifying mass shifted peptides and mapping crosslinking sites without further derivatization of the crosslinker rendered these additional studies unnecessary.

A limitation of the current studies is that the frequency of crosslinking events could not be evaluated as observed peak intensities in MS studies reflect the relative ionization of the species, not its concentration. Accurate quantification would allow discrimination of crosslinking frequency. This limitation is a common problem in MS-based discovery studies that preclude the use of isotopically enriched standards. Future studies plan to incorporate a laser-induced fluorescence microfluidic platform (Davic and Cascio, manuscript in preparation) to sensitively quantify derivatized peptides.

In summary, CX-MS identified both intra- and unique intersubunit crosslinked peptides in the resting state of the human GlyR. MS/MS data for all mass-shifted peptides further refined the sites of crosslinking and provided distance restraints. In all cases, MS/MS product ion scans of mass-shifted precursor ions identified a single site of crosslinking or refined the site of crosslinking to a few residues within the mass shifted tryptic fragment. The crosslinking sites from Cys^41^ identified in MS/MS studies identified many intrasubunit crosslinks and a single unique intersubunit crosslink. Of note, these CX-MS studies do not require large amounts of purified receptor, as picomoles of protein in a single SDS-PAGE gel is sufficient to identify dozens of crosslinks and thus may be capable of identifying networks of crosslinks that will aid in modeling/refining the structure of GlyR in its resting state. Additionally, these studies were conducted in reconstituted lipid vesicles, in the absence of detergents. CXMS studies can be performed in a state-dependent manner in the presence or absence of agonists, antagonists, or allosteric ligands with any desired lipid composition. Future work will apply CXMS to systematically introduced single cysteines in homopentameric a1 GlyR to generate a more comprehensive matrix of distance constraints in the resting state and other functional states of this paradigmatic receptor. This information can be used to update and refine the models of pLGICS in different conformational states. This experimental approach is particularly useful in identifying networks of constraints for flexible loops that may be poorly resolved or missing in studies of GlyR and its structural homologs. A similar approach using azi-cholesterol, a photoreactive derivative of cholesterol, has been utilized by our group to examine the lipid accessible regions of GlyR^51^, as well as the serotonin transporter^52^. As a result, these CXMS approaches offer a complementary approach to other high-resolution biophysical studies to sensitively and accurately map distance constraints for highly allosteric membrane proteins in a native environment.

## ACKNOWLEDGMENT

This work was supported by grants from the NIH (R21 MH098127) and a PA Dept. of Health CURE award. The purchase of the mass spectrometer was supported by the NSF (MRIDBI-0821401).

