## Supplemental Table 1 for "Site-specific Crosslinking Coupled with Mass Spectrometry as a Structural Tool in Studies of the Human α_1_ Glycine Receptor"

| SDS Band | Peptide | Δppm | Modifications |
|---|---|---|---|
| Monomeric | $^{3}$SATKPMSPSDFLDKLMGR$^{20}$ | -9.48,1.2, 1.1 | XL; XL OXD |
| | $^{34}$GPPVNVSCNIFINSFGSIAETTMDYRVNIFLR$^{65}$ | 0.58, 5.49 | XL(Na) |
| | $^{60}$VNIFLRQQWNDPR$^{72}$ | 2.11,-8.34,3.35 | XL; XL(Na);XL |
| | $^{105}$GAHFHEITTDNKLLR$^{119}$ | 4.67, 2.15 | XL(Na); XL(Na) OXD |
| | $^{120}$ISRNGNVLYSIR$^{131}$ | -3.23, -8.24 | XL |
| | $^{123}$NGNVLYSIRITLTLACPMDLK$^{143}$ | -9.94, -1.37, 7.52 | XL(K) ALK OXD ACRYL; XL 2ACRYL; XL ALK OXD |
| | $^{253}$VGLGITTVLTMTTQSSGSRASLPK$^{276}$ | 9.35, 8.33, 6.31 | XL(K) ACRYL; XL(Na) |
| | $^{272}$ASLPKVSYVK$^{281}$ | 2.93, 5.06, 2.99, 3.04 | XL(Na); XL(K) |
| | $^{393}$IGFPMAFLIFNMFYWIIYK$^{411}$ | -0.01 | XL(Na) OXD ACRYL |
| Oligomeric | $^{3}$SATKPMSPSDFLDKLMGR$^{20}$ | -6.15, -1.44, -5.71, -1.04 | XL; XL OXD |
| | $^{17}$LMGRTSGYDARIRPNFK$^{33}$ | 9.39, -4.16, 6.2 | XL(Na); XL |
| | $^{34}$GPPVNVSCNIFINSFGSIAETTMDYRVNIFLR$^{65}$ | 3.82 | XL(Na) |
| | $^{60}$VNIFLRQQWNDPR$^{72}$ | -6.83, -6.56, -4.37 | XL |
| | $^{105}$GAHFHEITTDNKLLR$^{119}$ | 3.58, 5.96, 3.34, 4.24 | XL(Na); XL(Na) OXD |
| | $^{123}$NGNVLYSIRITLTLACPMDLK$^{143}$ | -8.23, 7.46, -9.67 | XL OXD, XL(K) OXD ALK |
| | $^{194}$DLRYCTKHYNTGK$^{206}$ | 1.41, -9.45, 0.67, -3.48 | XL(Na) ALK; XL(K) ALK OXD ACRYL; XL |
| | $^{253}$VGLGITTVLTMTTQSSGSRASLPK$^{276}$ | 7.9, 6.47, 6.7 | XL(K) ACRYL |
| | $^{272}$ASLPKVSYVK$^{281}$ | 2.1, -2.33, 2.05, 1.29 | XL(Na); XL(K) |
| | $^{393}$IGFPMAFLIFNMFYWIIYKIVR$^{414}$ | 3.12,2.26, 7.9 | XL;XL 2(OXD) |
| | $^{412}$IVRREDVHNQ$^{421}$ | 7.38 | XL OXD; |

**SI Table 1.** Assigned precursor mass shifted tryptic peptides extracted from monomeric and oligomeric bands. All Δppm for each independent trial are provided.
Modifications: XL-crosslinker; Na-sodium adduct; K-potassium adduct, ACRYL - acrylamidation; OXD-oxidatio[illegible]

n
